# Integrating Patient Metadata and Genetic Pathogen Data: Advancing Pandemic Preparedness with a Multi-Parametric Simulator

**DOI:** 10.1101/2023.08.22.554132

**Authors:** Maxime Bonjean, Jérôme Ambroise, Francisco Orchard, Alexis Sentis, Julie Hurel, Jessica S Hayes, Máire A Connolly, Jean-Luc Gala

## Abstract

Training and practice are needed to handle an unusual crisis quickly, safely, and effectively. Functional and table-top exercises simulate anticipated CBRNe (Chemical, Biological, Radiological, Nuclear, and Explosive) and public health crises with complex scenarios based on realistic epidemiological, clinical, and biological data from affected populations. For this reason, the use of anonymized databases, such as those from ECDC or NCBI, are necessary to run meaningful exercises. Creating a training scenario requires connecting different datasets that characterise the population groups exposed to the simulated event. This involves interconnecting laboratory, epidemiological, and clinical data, alongside demographic information.

The sharing and connection of data among EU member states currently face shortcomings and insufficiencies due to a variety of factors including variations in data collection methods, standardisation practices, legal frameworks, privacy, and security regulations, as well as resource and infrastructure disparities.

During the H2020 project PANDEM-2 (Pandemic Preparedness and Response), we developed a multi-parametric training tool to artificially link together laboratory data and metadata. We used SARS-CoV-2 and ECDC and NCBI open-access databases to enhance pandemic preparedness.

We developed a comprehensive training procedure that encompasses guidelines, scenarios, and answers, all designed to assist users in effectively utilising the simulator.

Our tool empowers training managers and trainees to enhance existing datasets by generating additional variables through data-driven or random simulations. Furthermore, it facilitates the augmentation of a specific variable’s proportion within a given set, allowing for the customization of scenarios to achieve desired outcomes.

Our multi-parameter simulation tool is contained in the R package *Pandem2simulator*, available at https://github.com/maous1/Pandem2simulator. A Shiny application, developed to make the tool easy to use, is available at https://uclouvain-ctma.Shinyapps.io/Multi-parametricSimulator/. The tool runs in seconds despite using large data sets.

In conclusion, this multi-parametric training tool can simulate any crisis scenario, improving pandemic and CBRN preparedness and response. The simulator serves as a platform to develop methodology and graphical representations of future database-connected applications.

## 1. Introduction

High-throughput sequencing, commonly referred to as next generation sequencing (NGS), has transformed biology and medicine (1).

This revolutionary technique enables the analysis of an extensive array of whole genome sequences (WGS), some of which being available in repositories like NCBI (2) and GISAID (3). By harnessing the power of NGS, this breakthrough has enabled real-time monitoring of ongoing outbreaks by unravelling the genetic evolution of pathogens (4,5). This encompasses the natural selection, taxonomy, and prevalence of new variants (6), as well as assessments of pathogenicity and gain-of-function attributes such as transmissibility and immunity evasion advantages (7).

Integration of NGS data with contextual metadata is of utmost importance in a dynamic environment characterised by rapidly evolving infectious diseases. These data are essential for making informed public health decisions (8,9). Age, gender, comorbidities, vaccination history, and disease outcomes are just a few of the biological, clinical, and epidemiological details covered by this contextual metadata (*Figure 1*). Particularly noteworthy among the most significant components of this metadata constellation are the contagiousness, hospitalisation, and deaths (10).

**Figure 1.**
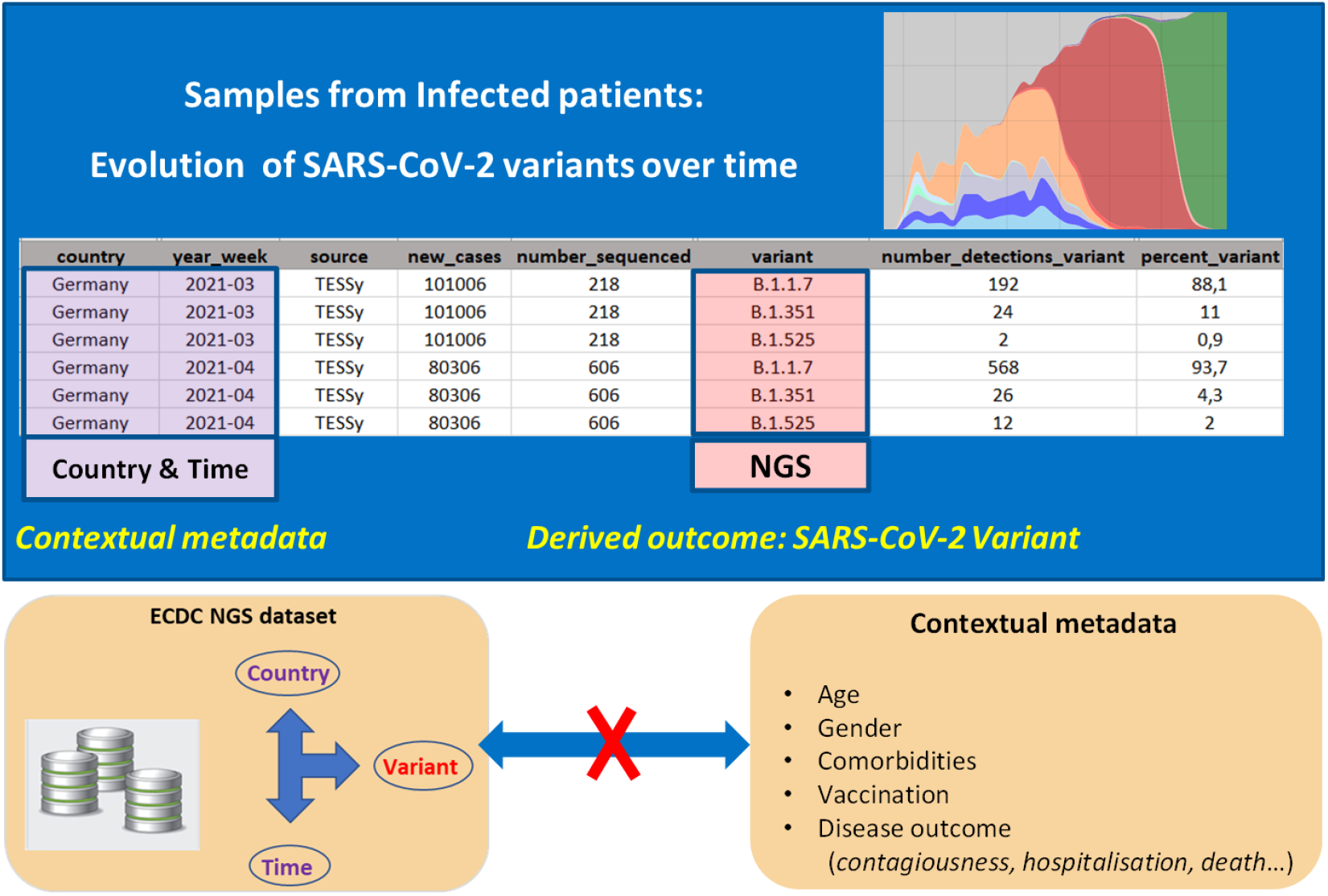
Illustration of the missing link between contextual metadata and Next Generation Sequencing (NGS) datasets. The figure shows weekly time series of NGS data provided by the TESSy dataset (source ECDC, https://www.ecdc.europa.eu/en/publications-data/data-virus-variants-covid-19-eueea) to monitor the evolution of SARS-CoV-2 variants in one EU Member State.

Pooling and interconnecting these data from various sources and sharing it across interdisciplinary teams allows for the identification of patterns, correlations, and emerging trends that would be difficult to discern from isolated datasets. Therefore, the ideal scenario would be for this integration to take place during data analysis in order to provide a real-time picture of an outbreak in terms of genetic evolution of the causative agent, its spatial and temporal dispersion and impact on efficacy of control measures and disease outcome (11).

However, there is a limited amount of open-access contextual metadata linked to pathogen sequencing data (*Figure 1*) (8). Sharing and interconnecting these data across countries and institutions currently face shortcomings and insufficiencies which stem from a variety of factors, including variations in data collection and standardisation methods, disparities in legal frameworks, and divergent privacy and security regulations, as well as resource and infrastructure gaps (12,13).

During the Horizon 2020 (H2020) project PANDEM-2 (Pandemic Preparedness and Response, https://pandem-2.eu/), we addressed these challenges in order to improve pandemic preparedness and response through a dedicated training platform. To achieve this objective, we developed a versatile multi-parametric training tool that simulates links between real or synthetic NGS data and contextual metadata. We used SARS-CoV-2, accessed ECDC and NCBI open-access databases, and developed the tool using an R coding package. This tool was then complemented by a user-friendly Shiny application to enhance accessibility and ease of use.

Our work involves assimilating data from ECDC’s COVID-19 weekly reports and NCBI, leading to the creation of time series datasets that encompass confirmed cases categorised by variant, country, and age group. Furthermore, we have integrated hospitalisation statistics stratified by variant, vaccination status, and country. As an added benefit, we have created a comprehensive training programme complete with guidelines, scenarios, and solutions to assist users in fully utilising the potential of our simulator.

This multi-parametric simulation tool represents a significant step forward in bridging the gap between NGS data and contextual metadata, paving the way for comprehensive and effective pandemic management. In practical terms, it does provide policymakers, public health officials, and health professionals with the opportunity to engage with a variety of scenarios and inputs through simulated synthetic data-based exercises, thereby improving evidence-based decision-making in pandemic response. Furthermore, the simulator serves as a platform to develop methodology and graphical representations of future database-connected applications.

In summary, this initiative is particularly crucial in light of current challenges, as it underpins our commitment to enhancing preparedness for major pandemics through the H2020 PANDEM-2 project.

## 2. Materials and methods

Our multiparameter simulation tool enables the user to change an existing initial dataset corresponding to the evolution of the number of cases during a specified period.

Accordingly, the initial dataset should include at least two variables: (i) the time expressed in a pre-defined format (e.g., YY-MM-DD) and (ii) the incidence expressed as an integer. It is worth noting that this incidence can correspond to the number of new cases or of new hospitalizations at any given time.

Aside from these two mandatory columns, the initial dataset can also include additional variables (e.g., localization [country name or region], vaccination status).

This initial dataset can be modified by the user through two distinct operations: (i) adding a new variable and (ii) modifying a relative risk characterising the association between two selected variables. These two operations can be used multiple times in order to add multiple distinct variables and/or modifying multiple distinct relative risks. The following steps are described in detail:

### 2.1 Methodological Steps

#### 2.1.1 Step 1: Addition of a new variable

The multi-parametric tool can be used to enrich the initial dataset (which represents the evolution of the number of infected cases) in two ways: A data-driven simulation or a random simulation (*Figure 2*).

a. *Data-driven simulation*: This method uses a KNN-based supervised classification method. The user must provide a second (named learning) dataset that has a variable (named X) that is also present in the initial dataset, as well as an additional variable (named Y) that will be added to the initial dataset. The KNN method learns the relationship between X and Y in the learning dataset and then predicts realistic values for this Y variable in the training dataset based on the values of the X variable. This straightforward method can be viewed as a means of connecting databases that were previously unconnected but share a common variable (for instance, time).
b. *Random simulation*: This is useful if the user wants to add another additional variable but does not have a learning database (as in data-driven simulation). The user can enter the name of the new categorical variable (for instance, vaccination status) as well as each of its levels (unvaccinated, first and second dose of vaccination). The user can then choose the desired appropriate proportion of each level.

**Figure 2.**
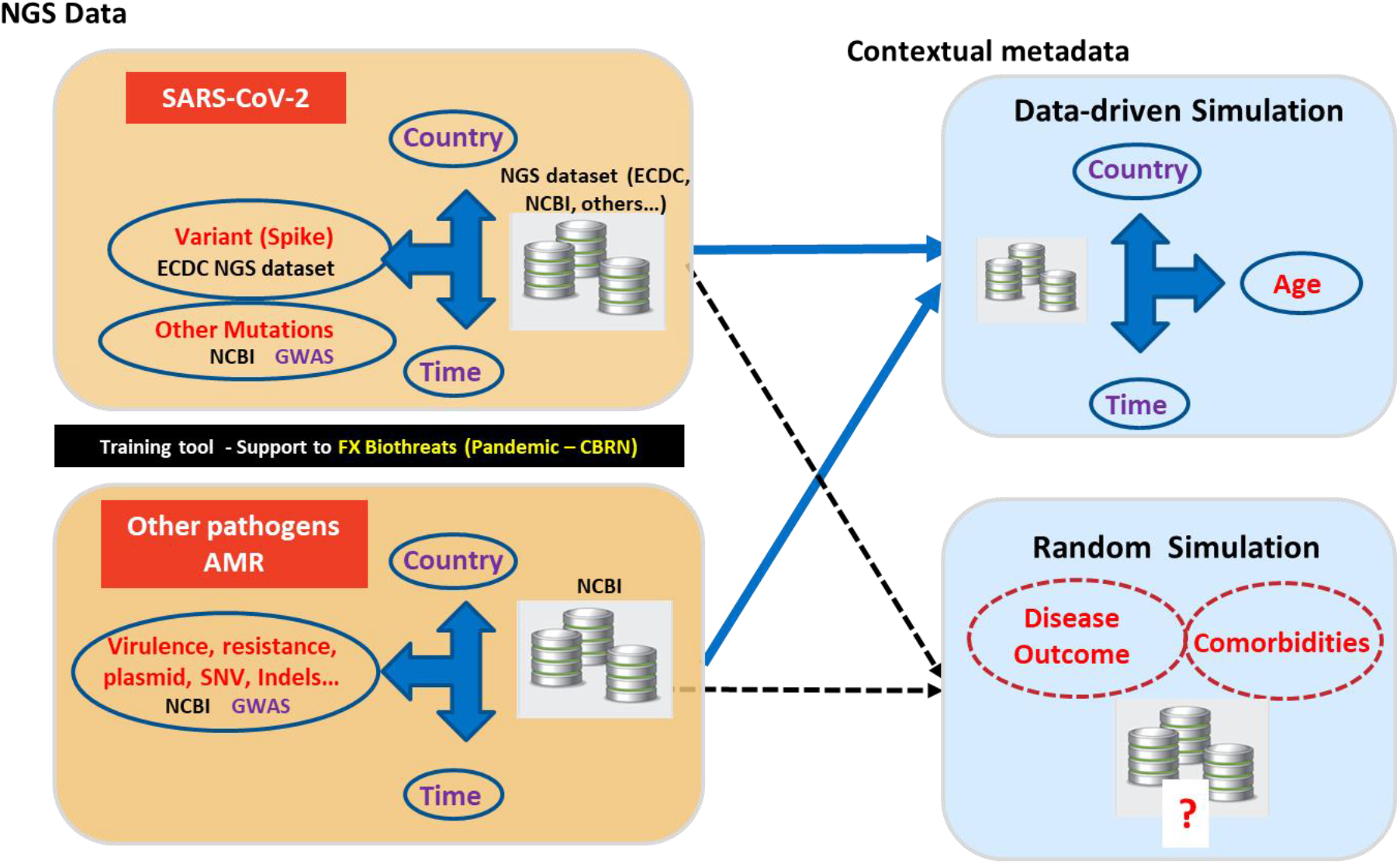
Data-driven and/or random simulations are employed to introduce an additional variable into the initial dataset using the multi-parametric simulation tool. These two types of simulation serve to enrich the initial dataset presented in the left panels, depicting distinct pandemic scenarios—SARS-CoV-2 in the upper panel and other pathogens (such as bacteria)/antimicrobial resistance (AMR) in the lower panel.

The upper left panel illustrates the SARS-CoV-2 time-series variant dataset provided by primary databases. The NGS dataset from ECDC and NCBI provides variables such as ‘variant’ in relation to ‘time’ and ‘country’. Information about other mutations and genome wide association studies (GWAS) can be found only in NCBI databases. By using common variables like ‘country’ and ‘time’, a data-driven simulation establishes a connection between the ‘variant’ variable and the contextual data as ‘age’ of the population group in which this variant is identified. if the contextual data ‘diseases outcome’ or ‘comorbidities’ are not accessible, linking them to the ‘variant’ variable would necessitate a random simulation.

The Lower left panel illustrates a time-series dataset representing the evolution of other pathogens, such as bacteria and antimicrobial resistance (AMR) genes. This data is sourced from the NCBI database. Using a similar approach as described above, both data-driven and random simulations are employed to establish connections between variables such as ‘virulence’, ‘resistance’, ‘presence of resistance plasmids’, ‘single nucleotide variant (SNV)’ or of ‘insertion-deletion (Indels)’.

#### 2.1.2 Step 2: Modifying a relative risk

In addition to enriching an initial dataset by adding new additional variables, the multivariate simulation tool also enables the user to manage the proportions of levels of variables in specific groups of patients (defined by specified levels of other variables) using the “enrichment” function T. The user sets the relative risk to a desired level. This function is therefore very useful as an epidemic training session because it enables the creation of different scenarios. It is worth noting that the RR is modified while maintaining the same proportions of the levels of each variable.

We developed our multi-parameter simulation tool using the R package, available at https://github.com/maous1/Pandem2simulator. We practised functional programming with the purrr package (14). To make the tool as user-friendly and ensure a widespread accessibility, we created a Shiny application: https://uclouvain-ctma.Shinyapps.io/MultiparametricSimulator/.

### 2.2 Use of the tool to enrich the COVID-19 training dataset (CD19TD)

The multi-parametric simulation tool was used to enrich the C19TD dataset created as part of the H2020 PANDEM-2 project (15). This C19TD database was developed in collaboration with several European public health and first responder agencies involved in the COVID-19 response, which helped define a list of variables and indicators to meet current and future pandemic management demands at the national and European level.

### 2.3 Training session to teach non-expert trainees how to use the multi-parametric tool

As part of H2020 project PANDEM-2 to develop a pandemic preparedness training platform, we conducted a training session described below. On February 1 2023, a training session was organised for 20 PANDEM-2 project participants. These participants included first responders, epidemiologists, public health experts, analysts, software developers, individuals from supply chain management, and disaster management professionals. During the training session, the trainees were shown how to use the multi-parametric tool for 30 minutes. This step was followed by a hands-on training activity in which they were tasked with creating datasets for the following pandemic scenarios:

- New SARS-CoV-2 variant of concern (VOC) propagating rapidly in young people: trainees were asked to generate a realistic dataset for a FX corresponding to the evolution of the number of infected cases by this new Sars-CoV-2 VOC.
- Highly virulent and resistant bacteria which spreads mainly in immunocompromised people and/or above 65 years old: trainees were asked to generate a dataset for a FX corresponding to the evolution of the number of infected cases.
- Influenza H1N1 virus with a risk of hospitalisation depending on vaccination status and age group: trainees were asked to generate a dataset for a FX corresponding to the evolution of the number of infected cases.

A questionnaire-based evaluation of the multiparametric simulator was presented to the trainees following the training session to assess the effectiveness of the simulation tool in generating data and facilitating functional exercises (Additional file 1: Questionnaire S1). The questions encompassed the clarity of training explanations concerning variable generation methods, data enrichment concepts, and dataset visualisation. The tool’s ability to modify and visualise datasets was examined, along with inquiries about usability improvements. Lastly, suggestions for tool development, layout refinement, and content enhancement were invited, encapsulating the assessment’s scope.

### 2.4 Use of the multi-parametric tool in a functional exercise (FX)

In the context of crisis management and preparedness, a FX is a type of simulation or drill used to test and improve the capabilities and coordination of stakeholders involved in responding to a major crisis or emergency situation. The goal of FX is to simulate a realistic crisis scenario, allowing participants to practise their roles, responsibilities, and decision-making processes in a safe setting (16), and evaluate their responses.

On March 15-16, 2023, the multi-parametric tool was used in a FX simulating an influenza pandemic triggered by transmission of a novel viral strain from birds to humans. The FX tested the response strategies put in place by two interacting Public Health Emergency Operation Centres in the national public health agencies of neighbouring countries (Germany and the Netherlands) and assessed the preparedness levels.

## 3. Results

### 3.1 Shiny Application overview

The Shiny application features 5 tabs that correspond to 5 stages of the process. The first tab is dedicated to importing the initial dataset import (*Figure 3*). In the second tab the user can seamlessly introduce new variables to the dataset. The third tab allows the user to visualise the dataset based on 3 variables. The time (i.e., the first variable) is consistently represented on the x-axis. Meanwhile, the levels of the second variable are displayed in separate panels, and the levels of the third variable are distinguished through different colours. In the fourth tab, the user can modify the relative risk, a change that is displayed in the fifth tab.

**Figure 3:**
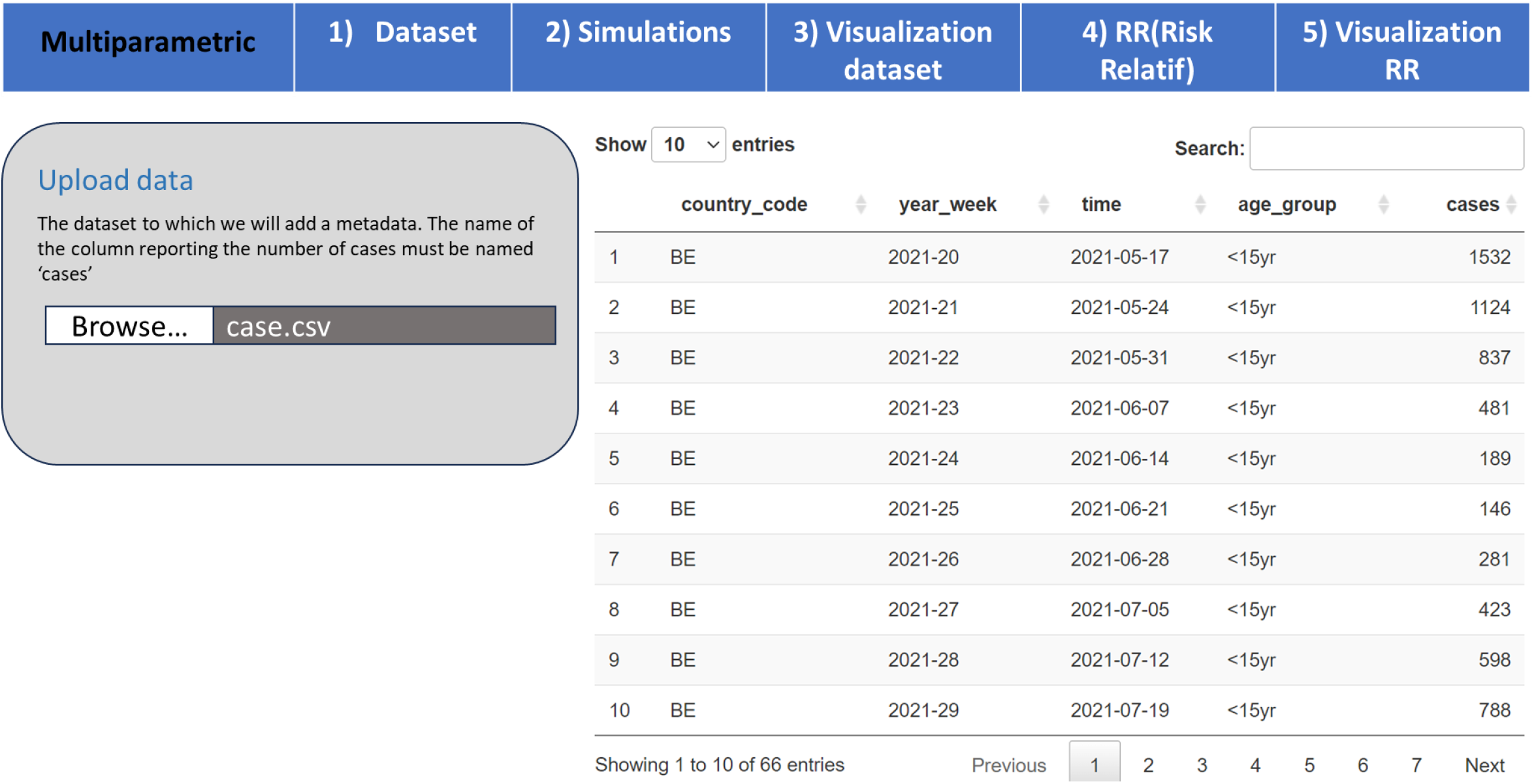
Screenshots of the First Tab in the Multi-Parametric Simulator Shiny Application.

### 3.2 Use of the tool to enrich the COVID-19 training dataset (CD19TD)

In addition, the multi-parametric simulation tool was used to enrich the C19TD dataset created as part of the H2020 PANDEM-2 project. This C19TD database was developed in collaboration with several European public health and first responder agencies involved in the COVID-19 response, who help define a list of variables and indicators to meet current and future pandemic management demands at the national and European level. For this application, data from ECDC’s COVID-19 weekly reports were used to generate a time series of confirmed cases by variant, country and age group, as well as hospitalisations by variant, vaccination status and country.

### 3.3 Training session

The training session with 20 participants demonstrated the user-friendliness and accessibility of the multi-parametric Shiny application. The evaluation provided by the trainees confirms that the concise video presentation of the tool and the subsequent creation of three distinct scenarios ensured a comprehensive understanding of its functionalities (*Table 1*).

**Table 1.**
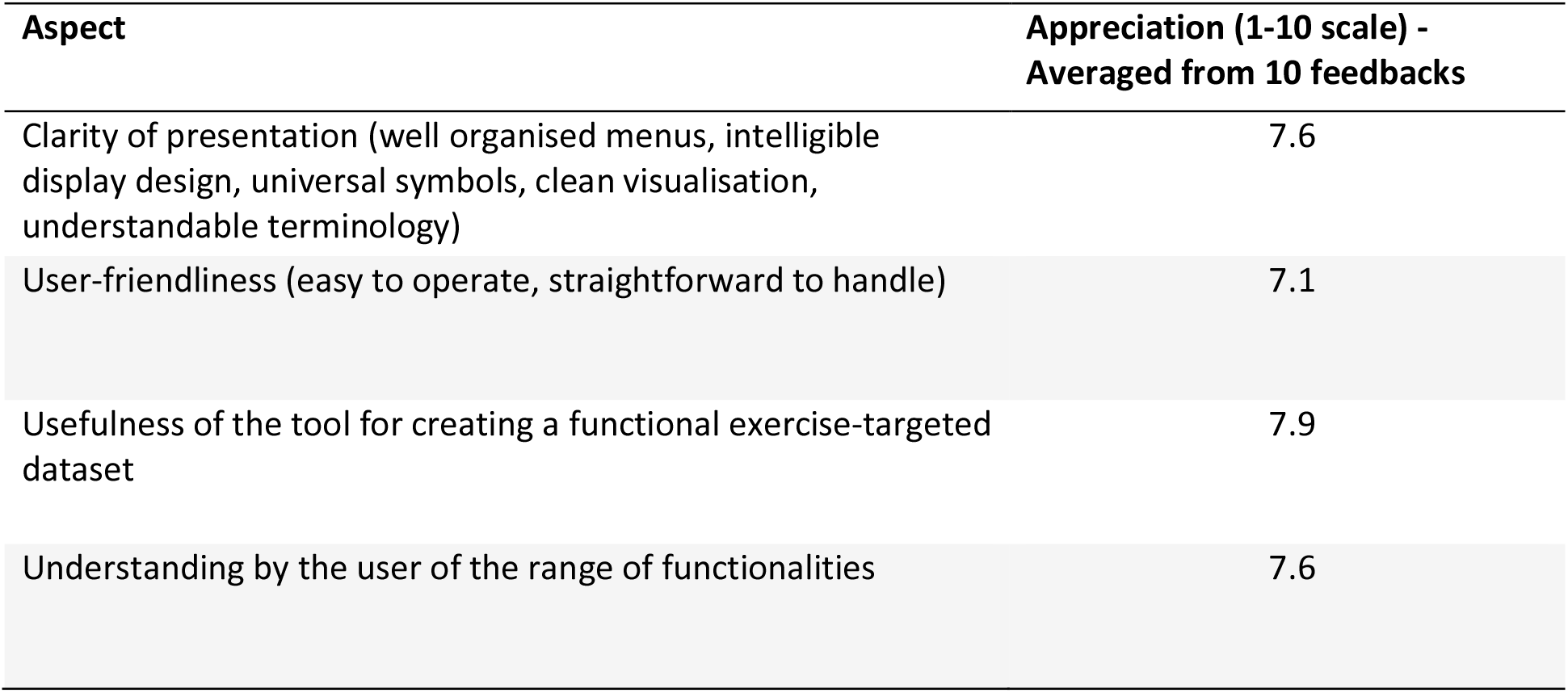
Overview of the training session participants’ evaluations.

| Aspect | Appreciation (1-10 scale) -<br>Averaged from 10 feedbacks |
| --- | --- |
| Clarity of presentation (well organised menus, intelligible display design, universal symbols, clean visualisation, understandable terminology) | 7.6 |
| User-friendliness (easy to operate, straightforward to handle) | 7.1 |
| Usefulness of the tool for creating a functional exercise-targeted dataset | 7.9 |
| Understanding by the user of the range of functionalities | 7.6 |

This successful training session indicates that the simulator can be effectively utilised by diverse stakeholders, including public health practitioners, researchers, and policymakers.

The initial datasets for these scenarios, along with solutions on how to modify them using the tool (Additional file 2: Dataset S1), and a video explaining the tool’s main functionalities are available as supplementary materials (Additional file 3: Video S1).

### 3.4 Application of the multi-parametric tool in a H2020 PANDEM-2 functional exercise (FX)

The FX organisers provided an initial dataset that included evolution of the number of cases over time. The multi-parametric tool was used to produce a numeric and graphic representation of the number cases over time. It was also used to add a new variable containing the name of the variant in the initial dataset. This new variable comprised two levels, which corresponded to the names of the previous (referred to in the FX as Alpha) and new (referred to in the FX as Gamma) strains that emerged in December according to the timeline of the pandemic scenario (*Figure 4*).

**Figure 4.**
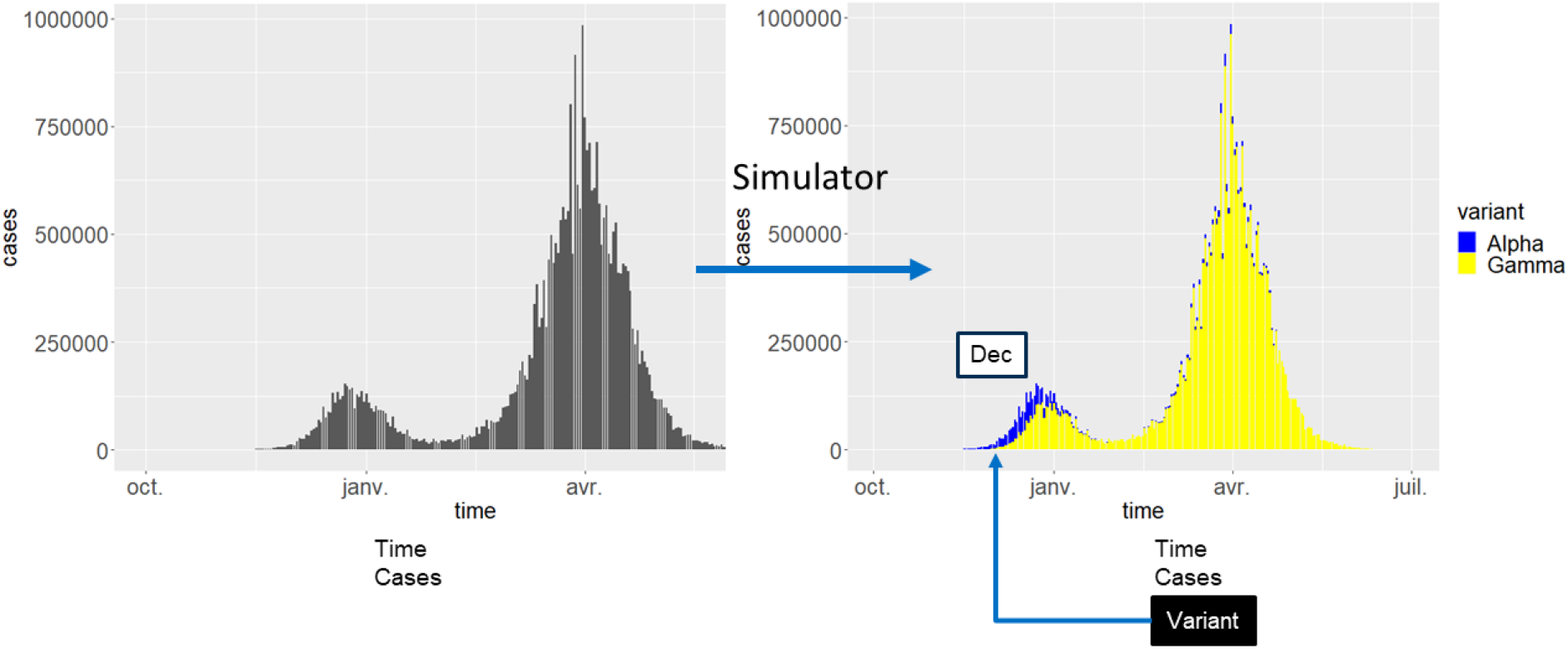
Use of the multi-parametric tool in a Functional exercise (FX). Simulation of the emergence of a Gamma variant of the H5N1 influenza virus during the course of the pandemic.

## 4. Discussion

The development and implementation of the multi-parametric simulation tool presented in this study marks a significant advancement in the field of preparedness and response to public health crises. The tool’s ability to enrich existing datasets and generate new variables through data-driven or random simulations offers valuable insights and practical applications in various crisis scenarios, including table-top and functional exercises (12). As stated before, enriching an initial dataset with a new variable by using the data-driven simulation can be viewed as a way to create links between databases that are initially not connected but have a common variable.

The FX conducted as part of the H2020 PANDEM-2 project demonstrated the effectiveness and adaptability of the simulation tool. The tool successfully demonstrated its ability to simulate the evolution of the number of cases during an influenza pandemic, as well as the emergence of a new strain on a specific date specified by the scenario. It is applicable to any type of infectious agent and its genetic evolution, in terms of strains or variants, depending on the specific scenario.

The flexibility of the simulation tool proved to be a valuable asset for evaluating response strategies implemented by two interacting Public Health Emergency Operation Centres and assessing their preparedness levels. The multi-parametric tool showcased its value as support to FX and for public health professionals in enhancing their crisis management skills.

Furthermore, the seamless flow of the training session, followed by its evaluation by the 20 participants, showcased the user-friendliness and accessibility of the multi-parametric Shiny application. The combination of a concise video presentation with a hands-on training session using the simulator to explore provided databases and generate data as required by the three proposed scenarios - focused on dynamic bacterial or viral infectious diseases - ensured a thorough understanding of the tool functionalities and capabilities. The success of this training session highlights the simulator’s effectiveness in educating a wide range of array of decision-makers including public health officials, health care professionals, researchers, and policymakers.

The integration of the multi-parametric simulation tool with the R programming language, as well as its encapsulation in the Pandem2simulator package, emphasises its potential for broader application and adoption within the scientific community. The functional programming approach, using the purrr (14) package, exemplifies the tool’s efficiency in managing large datasets while maintaining fast execution times, as demonstrated by the rapid execution of scenarios within a few seconds.

However, alongside significant achievements, certain limitations must be acknowledged. Notably, it is important to emphasise that, while the tool generates rich datasets, they are fictional and lack the same value as authentic data. The challenge lies in acquiring real metadata connected to pathogen genomic data and integrating them in anonymised databases. To facilitate data sharing between countries and institutions, future efforts should focus on addressing privacy, regulatory and standardisation concerns, as well as refining data management policies (17).

Moreover, continuous updates and improvements to the simulator’s functionalities and user interface would further enhance its usability and relevance. Nonetheless, the immersive learning experience provided by the simulator to key stakeholders in pandemic response significantly contributes to improve health emergency preparedness and response. In conclusion, the multi-parametric simulation tool developed as part of the H2020 PANDEM-2 project offers a powerful and versatile resource for policymakers, public health officials, and practitioners to prepare for and respond effectively to various crisis situations. By leveraging epidemiological, clinical, and biological metadata, the simulator is a first step in bridging the gap between pathogen genomic data and the integration of contextual metadata. Other pandemic simulation tools have been reported (18-21), but they address different aspects of pandemic requirements. Some concentrate on viral genealogy (18), while others on healthcare resource availability and the impact of shortages on public health (19), and still others on COVID-19 symptomatic case projections (21). The ability of our multi-parametric application to simulate a wide range of crisis scenarios, along with its adaptability for content and layout enhancements, makes it a valuable asset for generating the inputs needed for pandemics and CBRN FX. Given the continuous evolution of threats, sometimes at an extremely rapid pace, this simulation tool holds promise for future applications and contributions to global health security.

## Supporting information

Additional file 1: Questionnaire S1

## Author Contributions

Conceptualization, J.A., A.S., F.O. and J-L.G.; Methodology, J.A., MB., J.H, F.O., A.S. and J-L.G.; Validation, J.A., MB. and J-L.G.; Formal Analysis, J.A., M.B., F.O.; Investigation, J.A., MB. and J-L.G.; Resources, J-L.G.; Data Curation, J.A., M.B., J.H. and F.O.; Writing – Original Draft Preparation, M.B., J.A., and J-L.G.; Writing – Review & Editing, J.A., F.O., M.C. and J-L.G.; Visualization, J.A. and J-L.G.; Supervision, J-L.G.C.; Project Administration, J-L.G.; Funding Acquisition, J-L.G. and M.C.”

## Acknowledgments

We express our sincere gratitude to the entire H2020 PANDEM-2 technological team for their invaluable contributions to the development and implementation of the integrated dashboard featuring the multi-parametric simulator. Specifically, we would like to extend our appreciation to the following individuals for their dedicated efforts: C. Tighe, O; Zayed, C. Green, A. Ortiz, J. Albert, H. Chen (University of Galway, Ireland); G. Pacurar, B. Bucura, M. Enache (Clarisoft, Bucarest, Romania); C. Houareau (Robert Koch-Institut, Berlin, Germany); B. Kaluza, S. Römer (Fraunhofer, Munchen, Germany); L. Ngongalah (Trilateral Research LTD, London, United Kingdom).

## Data Availability Statement

The applications presented in this study are open-access. The link is provided in the manuscript.

