## Additional file 1: Questionnaire S1 for "Integrating Patient Metadata and Genetic Pathogen Data: Advancing Pandemic Preparedness with a Multi-Parametric Simulator"

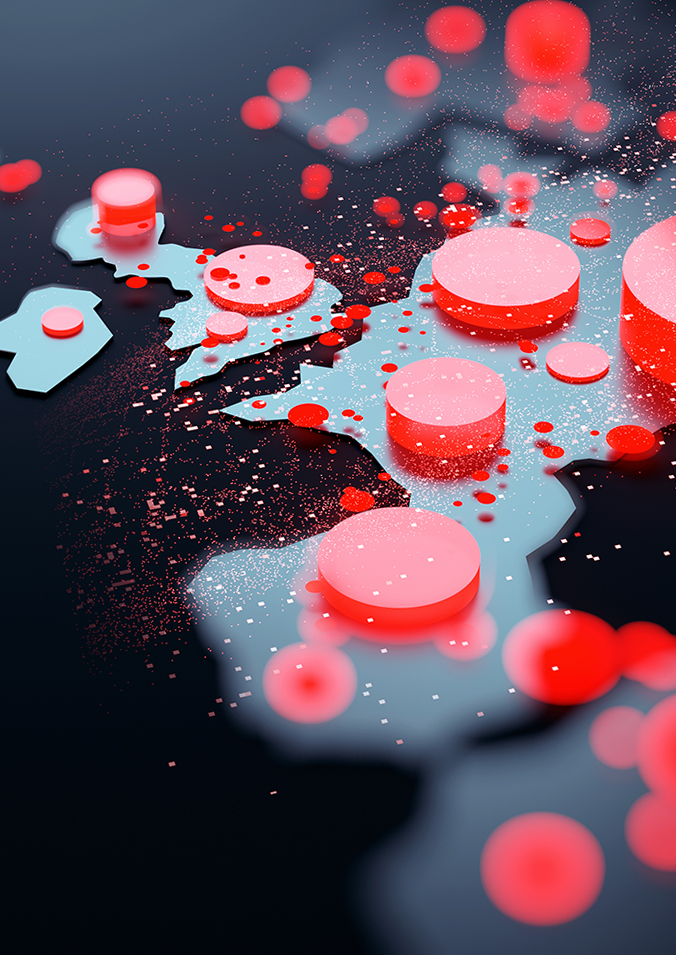

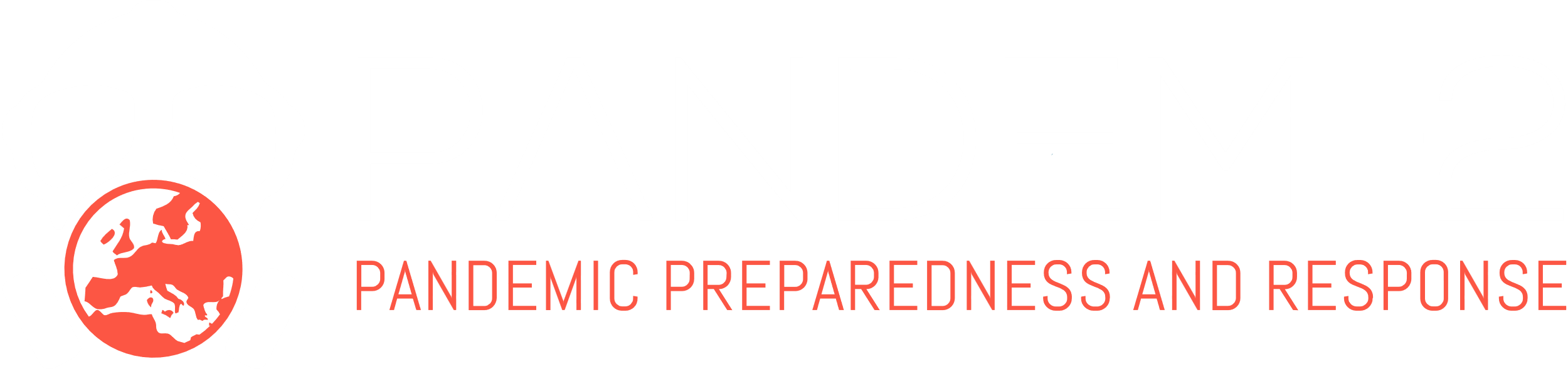

*Document submittal date*

Multi-parametric Simulator

The material presented and views expressed here are the responsibility of the author(s) only. The EU Commission takes no responsibility for any use made of the information set out.

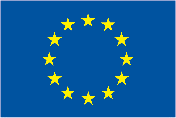

This project has received funding from the European Union’s Horizon 2020 research and innovation programme under Grant Agreement No. 883285

**Training Workshop**

### Training Workshop Evaluation

From perspective of the PANDEM-2 project, the training workshop has the goal of presenting the range of PANDEM-2 solutions to a wider audience of end-users to allow them to engage with the various systems, receive training in the presented functionalities. Secondly end-users are asked to give detailed evaluation feedback to rate how the systems score against the project key performance indicators (KPIs), as well as help improve the solutions further.

### Questions

These questions are adjusted to the respective training workshops “the Multi-parametric Simulator”

#### Introduction

The PANDEM-2 development team thanks you for participating in the PANDEM-2 IT tool training & evaluation workshop!

The trainings are designed to provide end-users an opportunity to understand the outcomes of the work done in the PANDEM-2 consortium over the past 24 months: you are invited to engage with the various IT tools created to support pandemic management, to receive training in the presented functionalities and to provide feedback on the tools to help defining a roadmap for future developments.

In order to best deliver on our mission: *providing novel solutions for EU pandemic preparedness and response*, we rely on your feedback as a participant, and kindly ask you to fill out the following questionnaire in full.

The survey is intended to be anonymous and all efforts have been made to ensure anonymity.

The PANDEM-2 project has received funding from the European Union’s Horizon 2020 research and innovation programme under grant agreement No 883285. The material presented and views expressed here are the responsibility of the author(s) only. The EU Commission takes no responsibility for any use made of the information set out.

#### Background & experience of user

1. How do you describe your professional role related to pandemic preparedness /management /response?
2. Are you working in a context, in which the Multi-parametric Simulator could be applied?

#### Training and documentation

1. Could you follow the workshop and was the training understandable? [Y/N]
   a. If not, please specify: []
2. Was the training material helpful and complete (presentations, documents, videos)? [Y/N]
   a. If not, please specify: []
3. Did you have all necessary information to engage with the Multi-parametric Simulator? [Y/N]
4. On a scale from 1 (not confident) to 10 (very confident),

how confident are you that you understood the range of functionalities of the Multi-parametric Simulator and how to apply them? [1-10]

5. Did you understand what was expected of you and fulfil the tasks given to the participants (if any)? [Y/N]
   a. If not, what could be improved? []
6. Will you easily remember how to use the Multi-parametric Simulator or could you use it again without written instructions? [Y/N]
   a. If not, please specify: []

#### Usability and user experience

*The usability and user experience of the PANDEM-2 solution is assessed by the end-users to be equal or higher than 7 out of 10****.***

1. Is the information in the the Multi-parametric Simulator clearly presented (well organized menus, intelligible display design, universal symbols, clean visualization, understandable terminology)?
   a. On a scale from 1 (poor presentation) to 10 (very well presented)? [1-10]
2. How would you evaluate the user-friendliness (easy to operate, straightforward to handle) of the Multi-parametric Simulator ?
   a. On a scale from 1 (not user-friendly) to 10 (very user-friendly)? [1-10]
3. Is the usage of the Multi-parametric Simulator easy to learn?
   a. On a scale from 1 (difficult & complicated) to 10 (self-explaining)? [1-10]
4. Do you have any recommendations to improve the usability of the Multi-parametric Simulator? [Y/N] always []

#### Perceived usefulness

*The perceived usefulness of the PANDEM-2 solution is assessed by the end-users to be equal or higher than 7 out of 10.*

1. On a scale of 1-10 (1: not helpful, 10: very helpful), please rate the possible usefulness of implementing the Multi-parametric Simulator in:
   a. overall pandemic planning and response? [1-10]
   b. your area of work? [1-10]
2. The multiparametric tools allow the generation of new variables using 2 distinct methods: ”random simulation” and ”data-driven simulation”. The latter method can generate a more realistic distribution of data than the former.
a. Do you think these two options were clearly explained during the training? [Y/N]
   i. If not, please specify: []
b. On a scale of 1-10 (1 being the least, 10 being the highest), do you think that both options can be useful in generating data for a functional exercise? [1-10]
3. The multiparametric tools enable data enrichment in order to modify an existing dataset. This step makes no changes to the number of cases or variables. Enrichment is the process of altering the proportion of a variable in a specific subgroup defined by one or more variables.
a. Do you think that this concept was clearly explained during the training? [Y/N]
   i. If not, please specify: []
b. On a scale of 1-10 (1 being the least, 10 being the highest), do you think that such a tool is useful for creating a functional exercise-targeted dataset? [1-10]
c. Is there another type of transformation/feature you would like to implement if you were using or creating a dataset for a functional exercise? []
4. The multiparametric tools allow the user to view and/or modify the dataset that has been created. Although this dataset may contain a large number of columns/variables (for example, 8), the user can only see a subset of these columns/variables. The x-and y-axis of each graph are always the “time” and ”number of cases (or hospitalizations)”, respectively. In addition, the user can specify which variable should be displayed in (i) different colours and (ii) different panels.
a. Do you think that this concept was clearly explained during the training? [Y/N]
   i. If not, please specify: []
b. What aspects of this visualisation function could or should be improved? []
5. Do you have any suggestions
a. to further develop the multiparametric simulator? []
b. to refine the layout? []
c. to improve the current content of this application? []

**Effectiveness of solution**

*End-users assessed the positive impact of the PANDEM-2 solutions on training pandemic planning and response in relevant areas (Surveillance & Epidemiology, Command, Control & Communication, Emergency Public Information & Warning, Surge Capacity, Disease Prevention & Control) to be equal or higher than 2 in each of these categories (on a scale from 1 – no impact to 5 – great impact).*

1. The WHO outlines a set of relevant areas, which are critical for pandemic planning and response. On a scale from 1 (no impact) to 5 (great impact), do you think the Multi-parametric Simulator can make a positive impact on:
   a. Surveillance & Epidemiology	[1-10]
   b. Command, Control & Communication	[1-10]
   c. Emergency Public Information & Warning	[1-10]
   d. Surge Capacity	[1-10]
   e. Disease Prevention & Control	[1-10]

**Platform compatibility**

*The compatibility of the PANDEM-2 solution (to plans, procedures and systems of organisations) is assessed by the end-users to be equal or higher than 7 out of 10.*

1. Did you identify any conflicts between the procedures of the Multi-parametric Simulator compared with the regular plans and procedures in your organization? [Y/N]
   a. If so, please specify: []
2. Do you expect problems, when using the Multi-parametric Simulator in interaction with your system? [Y/N]
   a. If so, please specify: []
3. On a scale of 1-10 (1 being the least, 10 being the highest),

how do you rate the compatibility of the PANDEM-2 solution(s) presented today to plans, procedures and systems used in your organization? [1-10]
