## Additional file 2: Dataset S1 for "Integrating Patient Metadata and Genetic Pathogen Data: Advancing Pandemic Preparedness with a Multi-Parametric Simulator": Solution-exercise_1.pdf

### Multiparametric simulator

# -

### Training Tool

# -

### Exercise n°1

Maxime Bonjean, Julie Hurel, Jérôme Ambroise
Jean-Luc Gala
Center for Applied Molecular Technologies (CTMA)
UCLouvain, BE

CTMA
Centre de Technologies Moléculaires Appliquées

UCLouvain

**Exercise n°1 :** Generate a realistic dataset for a FX corresponding to the evolution of the number of infected cases by a new Sars-CoV-2 VOC which propagates much more rapidly in young people

**On the basis of the datasets found in the directory (see Google Drive and shared documents), please generate a dataset characterised by:**

- 5 variables (time, number of cases, age_group, vaccination status [vaccinated: 80%, un-vaccinated: 20%], VOC [generated by a data-driven simulation with the provided dataset from ECDC]).

- A specific VOC (B1.617.2) propagates much more rapidly in young people (<15yr) with a relative risk of 4.

### Multiparametric simulator
# -
### Training Tool
# -
### Solution to exercise n°1

### Exercise n°1

Multiparametric simulator | 1) Datasets | 2) Simulations | 3) Visualization datasets | 4) Enrichment | 5) Visualization enrichment

#### Upload data

**The dataset to which we will add a metadata. The name of the column reporting the number of cases must be named 'cases'.**

Browse... agegroup-cases-ex3.csv

Upload complete

**Show** 10 **entries** **Search:**

| | time | age_group | cases |
|---|---|---|---|
| 1 | 2021-05-17 | <15yr | 3032 |
| 2 | 2021-05-24 | <15yr | 2224 |
| 3 | 2021-05-31 | <15yr | 1637 |
| 4 | 2021-06-07 | <15yr | 881 |
| 5 | 2021-06-14 | <15yr | 289 |
| 6 | 2021-06-21 | <15yr | 346 |
| 7 | 2021-06-28 | <15yr | 481 |
| 8 | 2021-07-05 | <15yr | 823 |
| 9 | 2021-07-12 | <15yr | 1098 |
| 10 | 2021-07-19 | <15yr | 1488 |

Showing 1 to 10 of 66 entries

Previous 1 2 3 4 5 6 7 Next

### Exercise n°1

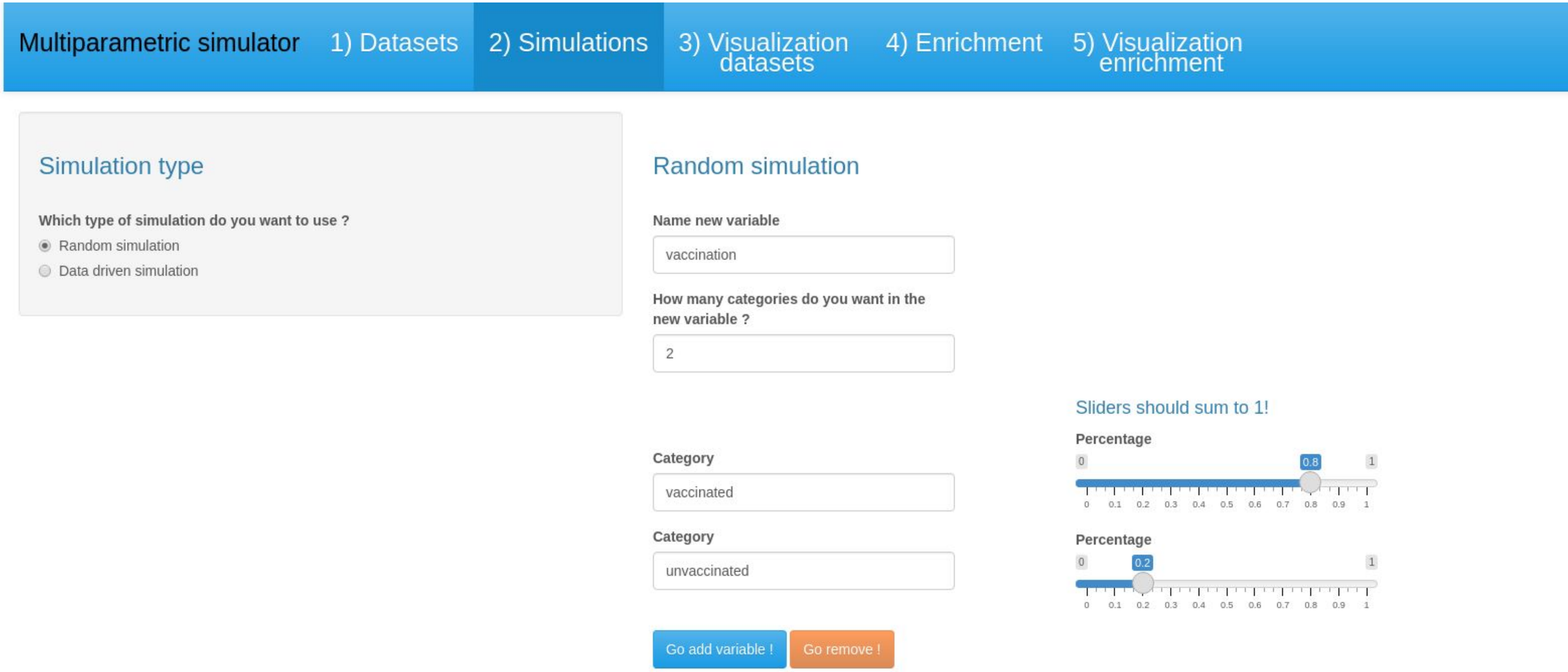
Multiparametric simulator
1) Datasets
2) Simulations
3) Visualization datasets
4) Enrichment
5) Visualization enrichment
Simulation type
Which type of simulation do you want to use ?
Random simulation
Data driven simulation
Random simulation
Name new variable
vaccination
How many categories do you want in the new variable ?
2
Category
vaccinated
Category
unvaccinated
Go add variable !
Go remove !
Sliders should sum to 1!
Percentage
0.8
Percentage
0.2

CTMA Centre de Technologies Moléculaires Appliquées

UCLouvain

### Exercise n°1

Multiparametric simulator | 1) Datasets | 2) Simulations | 3) Visualization datasets | 4) Enrichment | 5) Visualization enrichment

#### Simulation type

**Which type of simulation do you want to use ?**

- ○ Random simulation
- ◉ Data driven simulation

Add data | Advanced parameters

**Upload new dataset. The name of the column reporting the number of cases must be named 'cases'.**

Browse... variants.csv

Upload complete

**Select one column for outcome**

variant

**Select the names of the columns to match the data**

time

Go simulate! Go remove !

**Show** 10 **entries** **Search:**

| | time | age_group | vaccination | cases |
|---|---|---|---|---|
| 1 | 2021-05-17 | <15yr | unvaccinated | 306 |
| 2 | 2021-05-17 | <15yr | vaccinated | 1226 |
| 3 | 2021-05-17 | 15-24yr | unvaccinated | 613 |
| 4 | 2021-05-17 | 15-24yr | vaccinated | 2454 |
| 5 | 2021-05-17 | 25-49yr | unvaccinated | 1362 |
| 6 | 2021-05-17 | 25-49yr | vaccinated | 5446 |
| 7 | 2021-05-17 | 50-64yr | unvaccinated | 541 |
| 8 | 2021-05-17 | 50-64yr | vaccinated | 2164 |
| 9 | 2021-05-17 | 65-79yr | unvaccinated | 160 |

CTMA
Centre de Technologies Moléculaires Appliquées

UCLouvain

### Exercise n°1

Multiparametric simulator | 1) Datasets | 2) Simulations | 3) Visualization datasets | 4) Enrichment | 5) Visualization enrichment

#### Graph display of parameters

**Select the variable to be displayed in different colors (eg. variant)**

variant

**Select the variable to be displayed in different panels (eg. age_group)**

age_group

**Do you want to filter with a third variable?**

- Yes
- No

Display graph

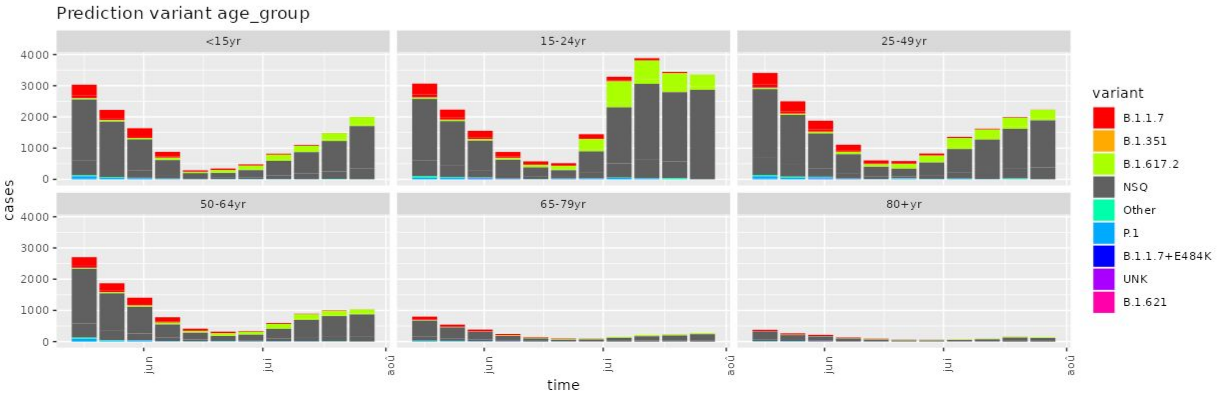

### Exercise n°1

Multiparametric simulator | 1) Datasets | 2) Simulations | 3) Visualization datasets | 4) Enrichment | 5) Visualization enrichment

#### Parameters

Specify the variable (e.g. variant) and the category (e.g. B.1.1.7) to enrich

Variable: variant

Category: B.1.617.2

Specify the variable(s) (e.g. age_group) and the category (e.g 25-49 yr) defining the population to be enriched

Variable: age_group

Category: <15yr

Add new variable and group

Display Relative Risk

Relative risk: 4

Go enrichment!

Show 10 entries

Search:

| | time | age_group | vaccination | variant | cases |
|---|---|---|---|---|---|
| 1 | 2021-05-17 | <15yr | unvaccinated | B.1.1.7 | 83 |
| 2 | 2021-05-17 | <15yr | unvaccinated | B.1.351 | 1 |
| 3 | 2021-05-17 | <15yr | unvaccinated | B.1.617.2 | 58 |
| 4 | 2021-05-17 | <15yr | unvaccinated | NSQ | 478 |
| 5 | 2021-05-17 | <15yr | unvaccinated | Other | 7 |
| 6 | 2021-05-17 | <15yr | unvaccinated | P.1 | 20 |
| 7 | 2021-05-17 | <15yr | vaccinated | B.1.1.7 | 330 |
| 8 | 2021-05-17 | <15yr | vaccinated | B.1.1.7+E484K | 5 |
| 9 | 2021-05-17 | <15yr | vaccinated | B.1.351 | 8 |
| 10 | 2021-05-17 | <15yr | vaccinated | B.1.617.2 | 84 |

Showing 1 to 10 of 654 entries

Previous 1 2 3 4 5 … 66 Next

Download .csv

CTMA
Centre de Technologies Moléculaires Appliquées

UCLouvain

### Exercise n°1

Multiparametric simulator | 1) Datasets | 2) Simulations | 3) Visualization datasets | 4) Enrichment | 5) Visualization enrichment

#### Graph display of parameters

**Select the variable to be displayed in different colors (eg. variant)**

variant

**Select the variable to be displayed in different panels (eg. age_group)**

age_group

**Do you want to filter with a third variable?**

- Yes
- No

Display graphs

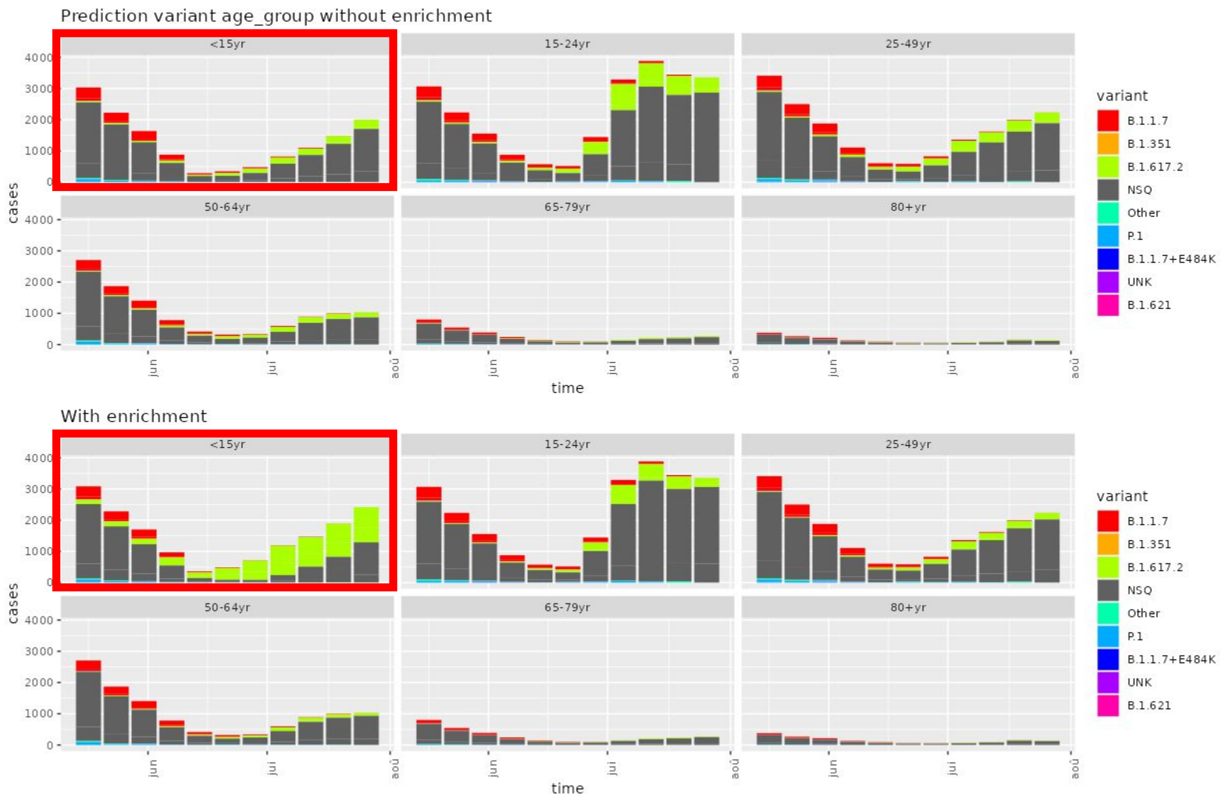
