## Additional file 2: Dataset S1 for "Integrating Patient Metadata and Genetic Pathogen Data: Advancing Pandemic Preparedness with a Multi-Parametric Simulator": Solution-exercise_2.pdf

### Multiparametric simulator
# -
### Training Tool
# -
### Exercise n°2

Maxime Bonjean, Julie Hurel, Jérôme Ambroise
Jean-Luc Gala
Center for Applied Molecular Technologies (CTMA)
UCLouvain, BE

CTMA
Centre de Technologies
Moléculaires Appliquées

UCLouvain

#### **Exercise n°2 :** Generate a dataset for a FX corresponding to the evolution of the number of infected cases by a highly virulent and resistant bacteria wich spreads mainly in immunocompromised people and/or above 80 YO

**On the basis of the datasets found in the directory (see Google Drive and shared documents), please generate a dataset characterised by:**

- 5 variables (time, number of cases, age_group, immunity status [normal: 80%, immune-compromised 20%], MLST [ST 69 50%, ST 515: 50%]).
- The ST 69 spreads more rapidly in older people (>80 yr) and/or those who are immunocompromised. The relative risk of spread in these categories is 6.

### Multiparametric simulator

# -

### Training Tool

# -

### Solution to exercise n°2

CTMA Centre de Technologies Moléculaires Appliquées

UCLouvain

### Exercise n°2

Multiparametric simulator | 1) Datasets | 2) Simulations | 3) Visualization datasets | 4) Enrichment | 5) Visualization enrichment

#### Upload data

**The dataset to which we will add a metadata. The name of the column reporting the number of cases must be named 'cases'.**

Browse... agegroup-cases.csv

Upload complete

**Show** 10 **entries** **Search:**

| | time | age_group | cases |
|---|---|---|---|
| 1 | 2021-05-17 | <15yr | 1532 |
| 2 | 2021-05-24 | <15yr | 1124 |
| 3 | 2021-05-31 | <15yr | 837 |
| 4 | 2021-06-07 | <15yr | 481 |
| 5 | 2021-06-14 | <15yr | 189 |
| 6 | 2021-06-21 | <15yr | 146 |
| 7 | 2021-06-28 | <15yr | 281 |
| 8 | 2021-07-05 | <15yr | 423 |
| 9 | 2021-07-12 | <15yr | 598 |
| 10 | 2021-07-19 | <15yr | 788 |

Showing 1 to 10 of 66 entries

Previous 1 2 3 4 5 6 7 Next

### Exercise n°2

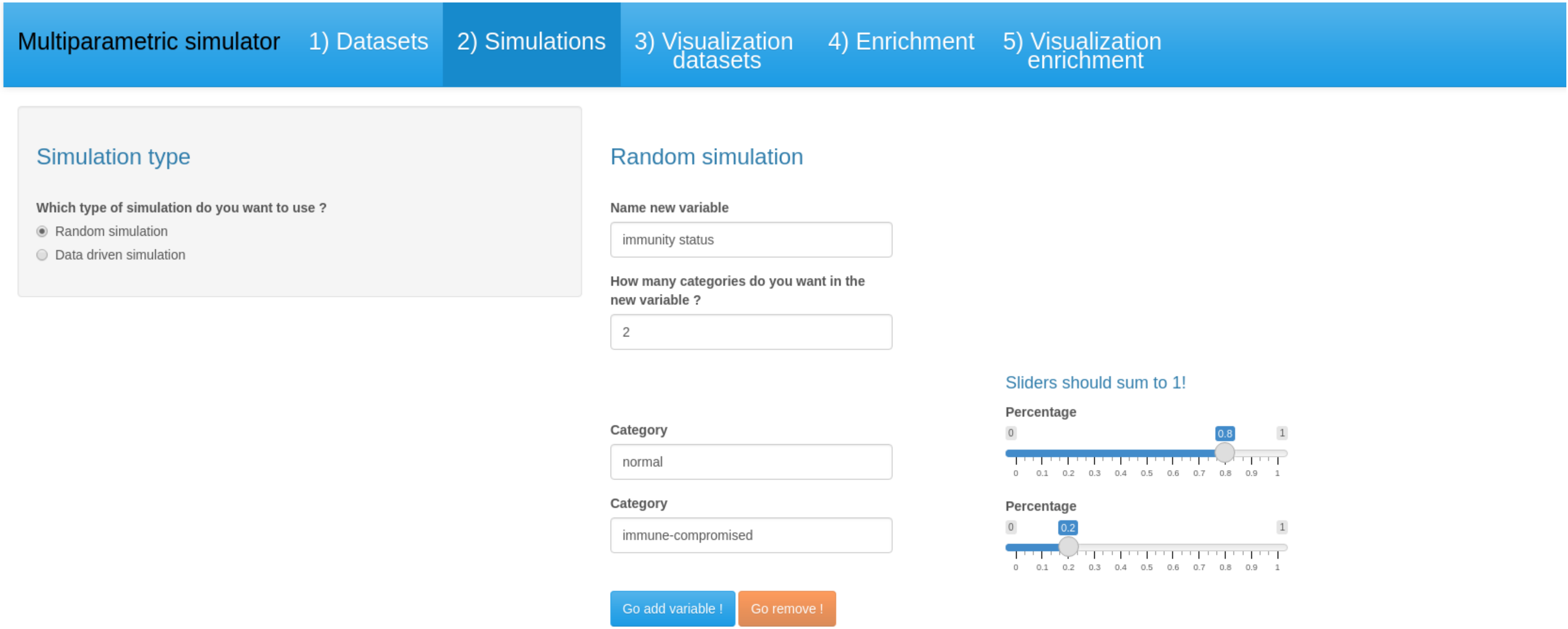
Multiparametric simulator
1) Datasets
2) Simulations
3) Visualization datasets
4) Enrichment
5) Visualization enrichment
Simulation type
Which type of simulation do you want to use ?
Random simulation
Data driven simulation
Random simulation
Name new variable
immunity status
How many categories do you want in the new variable ?
2
Category
normal
Category
immune-compromised
Go add variable !
Go remove !
Sliders should sum to 1!
Percentage
0.8
Percentage
0.2

CTMA
Centre de Technologies Moléculaires Appliquées

UCLouvain

### Exercise n°2

Multiparametric simulator | 1) Datasets | 2) Simulations | 3) Visualization datasets | 4) Enrichment | 5) Visualization enrichment

#### Simulation type

**Which type of simulation do you want to use ?**

- (•) Random simulation
- ( ) Data driven simulation

#### Random simulation

**Name new variable**

MLST

**How many categories do you want in the new variable ?**

2

**Sliders should sum to 1!**

**Category**

ST 69

**Percentage**: 0.5 (0 – 1)

**Category**

ST 515

**Percentage**: 0.5 (0 – 1)

Go add variable ! | Go remove !

**Show** 10 **entries** **Search:**

| | time | age_group | immunity_status | cases |
|---|---|---|---|---|
| 1 | 2021-05-17 | <15yr | normal | 1226 |
| 2 | 2021-05-17 | <15yr | immune-compromised | 306 |
| 3 | 2021-05-24 | <15yr | normal | 899 |
| 4 | 2021-05-24 | <15yr | immune-compromised | 225 |
| 5 | 2021-05-31 | <15yr | normal | 670 |
| 6 | 2021-05-31 | <15yr | immune-compromised | 167 |
| 7 | 2021-06-07 | <15yr | normal | 385 |

### Exercise n°2

**Multiparametric simulator** | 1) Datasets | 2) Simulations | 3) Visualization datasets | 4) Enrichment | 5) Visualization enrichment

#### Graph display of parameters

**Select the variable to be displayed in different colors (eg. variant)**

MLST

**Select the variable to be displayed in different panels (eg. age_group)**

age_group

**Do you want to filter with a third variable?**

- ○ Yes
- ◉ No

Display graph

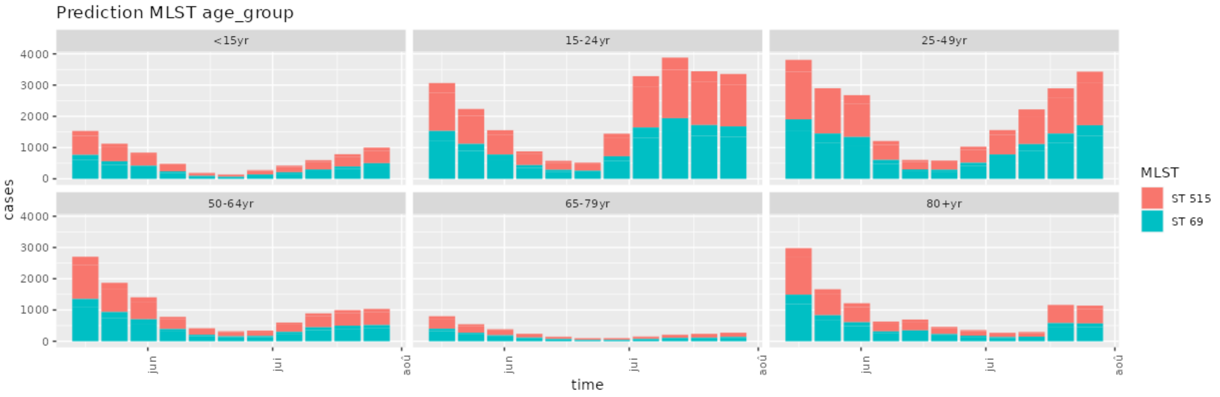

CTMA
Centre de Technologies Moléculaires Appliquées

UCLouvain

### Exercise n°2

Multiparametric simulator | 1) Datasets | 2) Simulations | 3) Visualization datasets | 4) Enrichment | 5) Visualization enrichment

#### Parameters

**Specify the variable (e.g. variant) and the category (e.g. B.1.1.7) to enrich**

**Variable**: MLST
**Category**: ST 69

**Specify the variable(s) (e.g. age_group) and the category (e.g 25-49 yr) defining the population to be enriched**

**Variable**: age_group
**Category**: 80+yr

**Variable**: immunity_status
**Category**: immune-compromised

× Click twice

Add new variable and group

Display Relative Risk

**Relative risk**

6

Go enrichment!

**Show** 10 **entries** **Search:**

| | time | age_group | immunity_status | MLST | cases |
|---|---|---|---|---|---|
| 1 | 2021-05-17 | <15yr | immune-compromised | ST 515 | 159 |
| 2 | 2021-05-17 | <15yr | immune-compromised | ST 69 | 147 |
| 3 | 2021-05-17 | <15yr | normal | ST 515 | 639 |
| 4 | 2021-05-17 | <15yr | normal | ST 69 | 587 |
| 5 | 2021-05-17 | 15-24yr | immune-compromised | ST 515 | 319 |
| 6 | 2021-05-17 | 15-24yr | immune-compromised | ST 69 | 293 |
| 7 | 2021-05-17 | 15-24yr | normal | ST 515 | 1278 |
| 8 | 2021-05-17 | 15-24yr | normal | ST 69 | 1176 |
| 9 | 2021-05-17 | 25-49yr | immune-compromised | ST 515 | 397 |
| 10 | 2021-05-17 | 25-49yr | immune-compromised | ST 69 | 365 |

Showing 1 to 10 of 264 entries

Previous 1 2 3 4 5 … 27 Next

Download .csv

### Exercise n°2

Multiparametric simulator | 1) Datasets | 2) Simulations | 3) Visualization datasets | 4) Enrichment | 5) Visualization enrichment

#### Graph display of parameters

**Select the variable to be displayed in different colors (eg. variant)**

MLST

**Select the variable to be displayed in different panels (eg. age_group)**

age_group

**Do you want to filter with a third variable?**

- Yes
- No

**Which one ?**

**Variable**

immunity_status

**Category**

immune-compromised

Display graphs

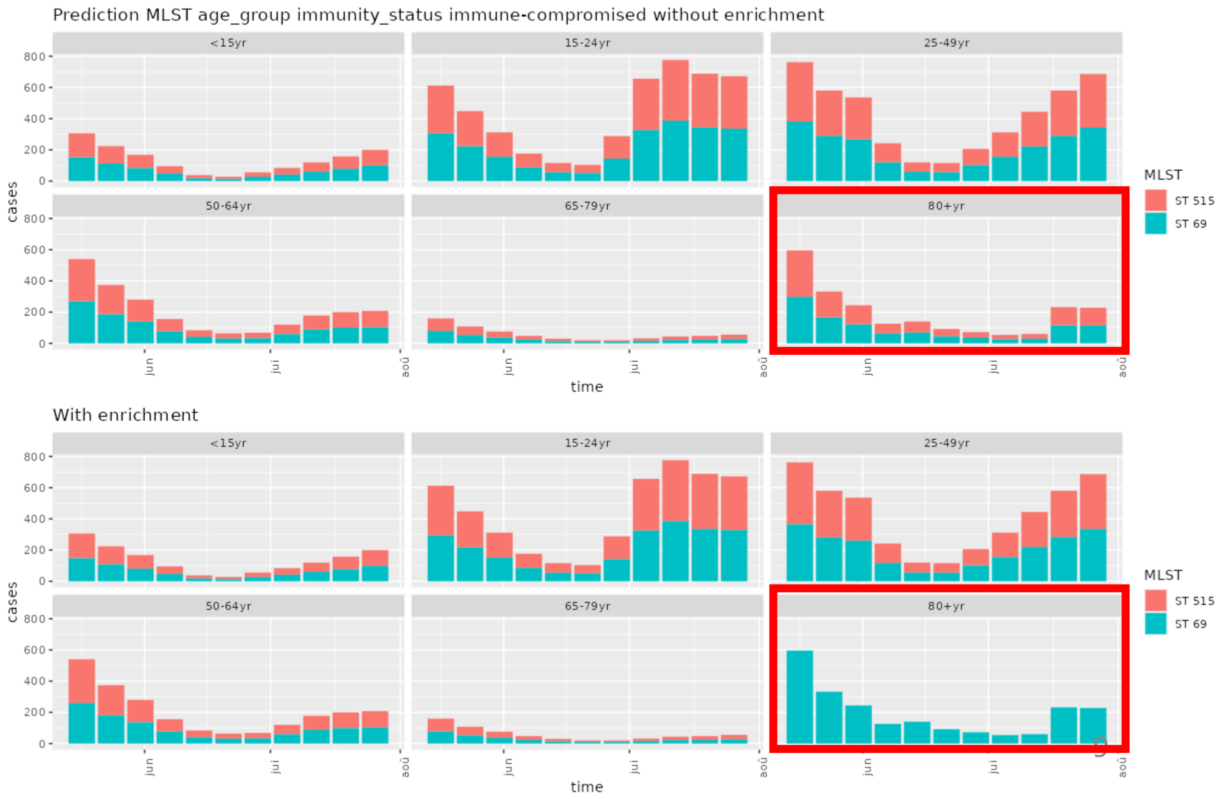
