## Additional file 2: Dataset S1 for "Integrating Patient Metadata and Genetic Pathogen Data: Advancing Pandemic Preparedness with a Multi-Parametric Simulator": Solution-exercise_3.pdf

### Multiparametric simulator

# -

### Training Tool

# -

### Exercise n°3

Maxime Bonjean, Julie Hurel, Jérôme Ambroise
Jean-Luc Gala
Center for Applied Molecular Technologies (CTMA)
UCLouvain, BE

#### **Exercise n°3 :** Generate a dataset for a FX corresponding to the evolution of the number of cases infected by influenza H1N1 virus with a risk of hospitalisation depending on vaccination status and age group

**On the basis of the datasets found in the directory (see Google Drive and shared documents), please generate a dataset characterised by:**

- 5 variables (time, number of cases, age_group, vaccination status [vaccinated: 50%, un-vaccinated: 50%], disease outcome [light symptoms 80%, hospitalisation 20%]).

- The disease outcome is affected by age_group (with a relative risk of hospitalisation of 3 for young patient <15 yr) and highly affected by vaccination status (with a relative risk of hospitalisation of 8 in unvaccinated patients).

### Multiparametric simulator
# -
### Training Tool
# -
### Solution to exercise n°3

CTMA
Centre de Technologies
Moléculaires Appliquées

UCLouvain

### Exercise n°3

Multiparametric simulator | 1) Datasets | 2) Simulations | 3) Visualization datasets | 4) Enrichment | 5) Visualization enrichment

#### Upload data

**The dataset to which we will add a metadata. The name of the column reporting the number of cases must be named 'cases'.**

Browse... agegroup-cases-ex3.csv

Upload complete

**Show** 10 **entries**

**Search:**

| | time | age_group | cases |
|---|---|---|---|
| 1 | 2021-05-17 | <15yr | 3032 |
| 2 | 2021-05-24 | <15yr | 2224 |
| 3 | 2021-05-31 | <15yr | 1637 |
| 4 | 2021-06-07 | <15yr | 881 |
| 5 | 2021-06-14 | <15yr | 289 |
| 6 | 2021-06-21 | <15yr | 346 |
| 7 | 2021-06-28 | <15yr | 481 |
| 8 | 2021-07-05 | <15yr | 823 |
| 9 | 2021-07-12 | <15yr | 1098 |
| 10 | 2021-07-19 | <15yr | 1488 |

Showing 1 to 10 of 66 entries

Previous 1 2 3 4 5 6 7 Next

### Exercise n°3

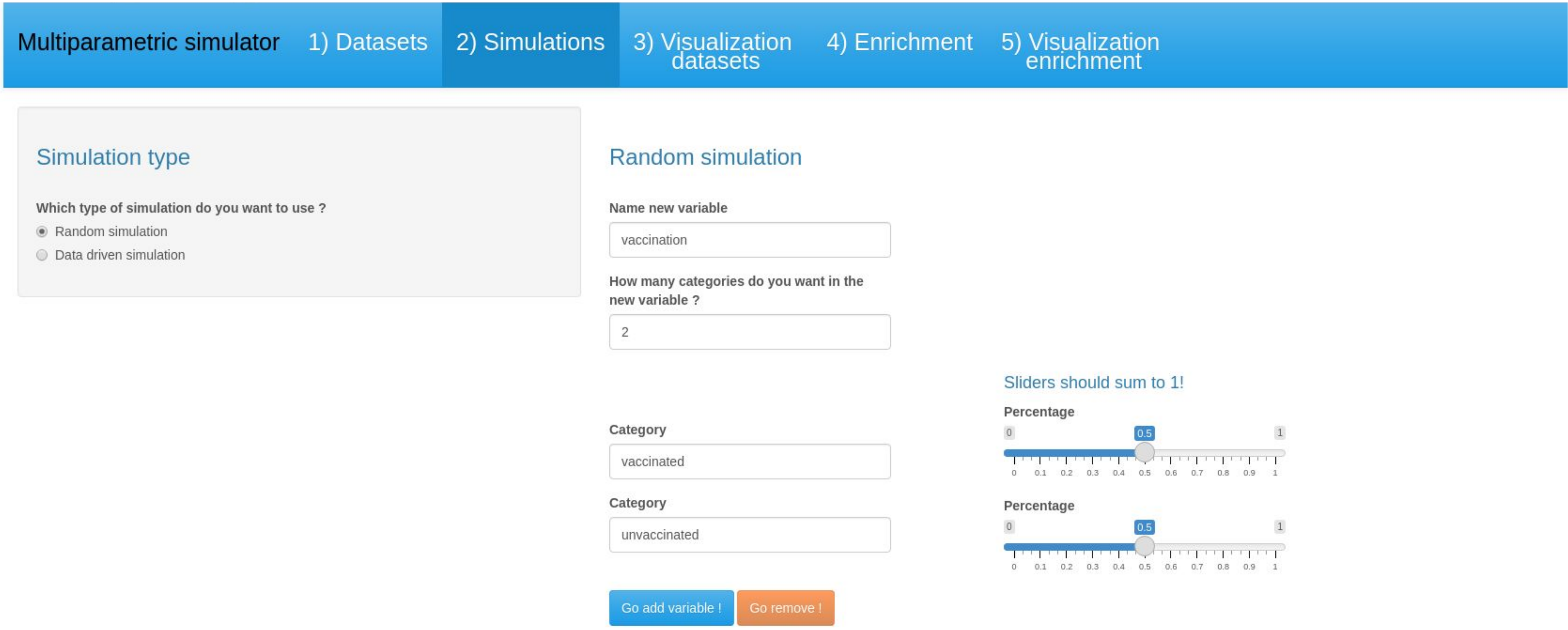
Multiparametric simulator
1) Datasets
2) Simulations
3) Visualization datasets
4) Enrichment
5) Visualization enrichment
Simulation type
Which type of simulation do you want to use ?
Random simulation
Data driven simulation
Random simulation
Name new variable
vaccination
How many categories do you want in the new variable ?
2
Sliders should sum to 1!
Category
vaccinated
Percentage
0.5
Category
unvaccinated
Percentage
0.5
Go add variable !
Go remove !

CTMA
Centre de Technologies Moléculaires Appliquées

UCLouvain

### Exercise n°3

Multiparametric simulator | 1) Datasets | 2) Simulations | 3) Visualization datasets | 4) Enrichment | 5) Visualization enrichment

#### Simulation type

**Which type of simulation do you want to use ?**

- (•) Random simulation
- ( ) Data driven simulation

#### Random simulation

**Name new variable**

disease outcome

**How many categories do you want in the new variable ?**

2

**Category**

light symptoms

**Category**

hospitalisation

Sliders should sum to 1!

**Percentage**

0 — 0.8 — 1

0 0.1 0.2 0.3 0.4 0.5 0.6 0.7 0.8 0.9 1

**Percentage**

0 — 0.2 — 1

0 0.1 0.2 0.3 0.4 0.5 0.6 0.7 0.8 0.9 1

Go add variable ! | Go remove !

**Show** 10 **entries**

**Search:**

| | time | age_group | vaccination | cases |
|---|---|---|---|---|
| 1 | 2021-05-17 | <15yr | vaccinated | 1516 |
| 2 | 2021-05-17 | <15yr | unvaccinated | 1516 |
| 3 | 2021-05-24 | <15yr | vaccinated | 1112 |
| 4 | 2021-05-24 | <15yr | unvaccinated | 1112 |
| 5 | 2021-05-31 | <15yr | vaccinated | 818 |
| 6 | 2021-05-31 | <15yr | unvaccinated | 818 |
| 7 | 2021-06-07 | <15yr | vaccinated | 440 |

### Exercise n°3

Multiparametric simulator | 1) Datasets | 2) Simulations | 3) Visualization datasets | 4) Enrichment | 5) Visualization enrichment

##### Graph display of parameters

**Select the variable to be displayed in different colors (eg. variant)**

disease_outcome

**Select the variable to be displayed in different panels (eg. age_group)**

age_group

**Do you want to filter with a third variable?**

- Yes
- No

Display graph

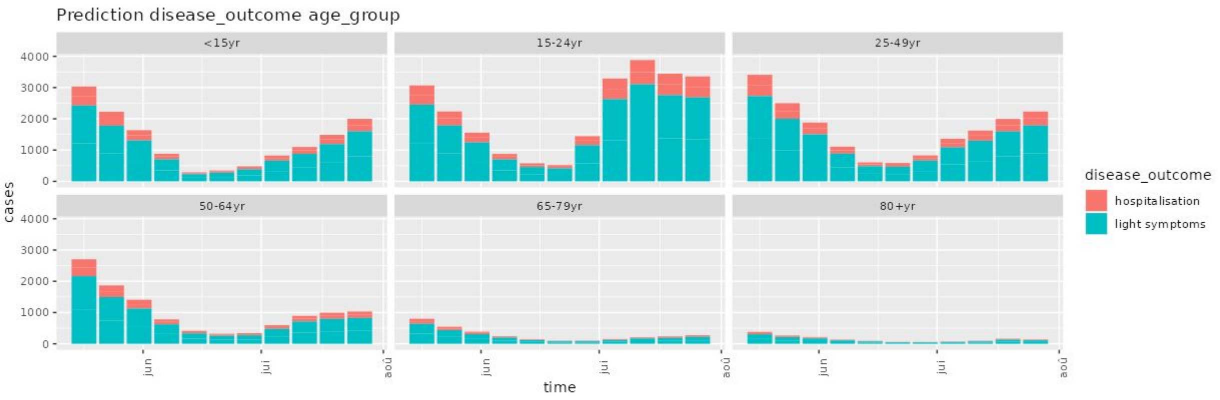

CTMA Centre de Technologies Moléculaires Appliquées

UCLouvain

### Exercise n°3

Multiparametric simulator | 1) Datasets | 2) Simulations | 3) Visualization datasets | 4) Enrichment | 5) Visualization enrichment

#### Parameters

Specify the variable (e.g. variant) and the category (e.g. B.1.1.7) to enrich

Variable: disease_outcome

Category: hospitalisation

Specify the variable(s) (e.g. age_group) and the category (e.g 25-49 yr) defining the population to be enriched

Variable: age_group

Category: <15yr

Add new variable and group

Display Relative Risk

Relative risk

3

Go enrichment!

Show 10 entries

Search:

| | time | age_group | vaccination | disease_outcome | cases |
|---|---|---|---|---|---|
| 1 | 2021-05-17 | <15yr | unvaccinated | hospitalisation | 948 |
| 2 | 2021-05-17 | <15yr | unvaccinated | light symptoms | 891 |
| 3 | 2021-05-17 | <15yr | vaccinated | hospitalisation | 948 |
| 4 | 2021-05-17 | <15yr | vaccinated | light symptoms | 890 |
| 5 | 2021-05-17 | 15-24yr | unvaccinated | hospitalisation | 212 |
| 6 | 2021-05-17 | 15-24yr | unvaccinated | light symptoms | 1322 |
| 7 | 2021-05-17 | 15-24yr | vaccinated | hospitalisation | 211 |
| 8 | 2021-05-17 | 15-24yr | vaccinated | light symptoms | 1323 |
| 9 | 2021-05-17 | 25-49yr | unvaccinated | hospitalisation | 235 |
| 10 | 2021-05-17 | 25-49yr | unvaccinated | light symptoms | 1469 |

Showing 1 to 10 of 264 entries

Previous 1 2 3 4 5 … 27 Next

Download .csv

CTMA
Centre de Technologies Moléculaires Appliquées

UCLouvain

### Exercise n°3

Multiparametric simulator | 1) Datasets | 2) Simulations | 3) Visualization datasets | 4) Enrichment | 5) Visualization enrichment

#### Graph display of parameters

**Select the variable to be displayed in different colors (eg. variant)**

disease_outcome

**Select the variable to be displayed in different panels (eg. age_group)**

age_group

**Do you want to filter with a third variable?**

- Yes
- No

Display graphs

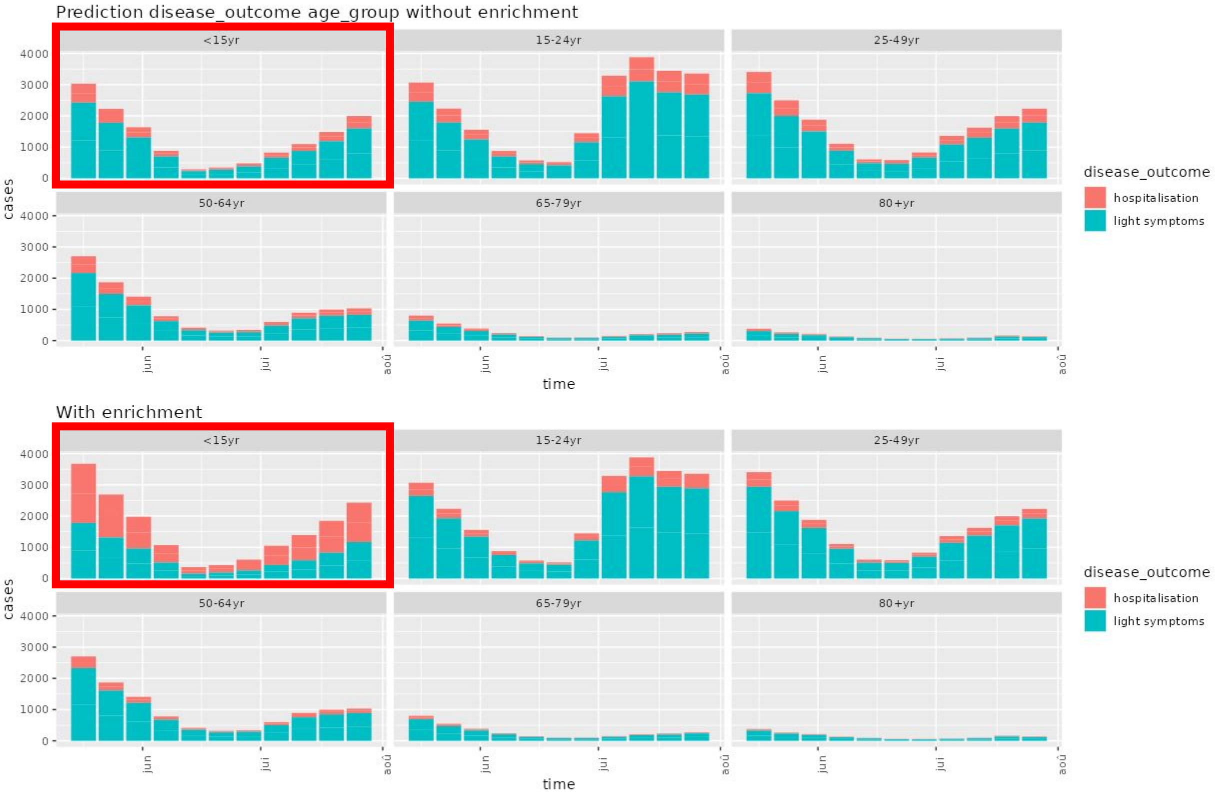

### Exercise n°3

Multiparametric simulator | 1) Datasets | 2) Simulations | 3) Visualization datasets | 4) Enrichment | 5) Visualization enrichment

#### Parameters

**Specify the variable (e.g. variant) and the category (e.g. B.1.1.7) to enrich**

**Variable**: disease_outcome
**Category**: hospitalisation

**Specify the variable(s) (e.g. age_group) and the category (e.g 25-49 yr) defining the population to be enriched**

**Variable**: vaccination
**Category**: unvaccinated

Add new variable and group

Display Relative Risk

**Relative risk**

8

Go enrichment!

Show 10 entries    Search:

| | time | age_group | vaccination | disease_outcome | cases |
|---|---|---|---|---|---|
| 1 | 2021-05-17 | <15yr | unvaccinated | hospitalisation | 1344 |
| 2 | 2021-05-17 | <15yr | unvaccinated | light symptoms | 977 |
| 3 | 2021-05-17 | <15yr | vaccinated | hospitalisation | 67 |
| 4 | 2021-05-17 | <15yr | vaccinated | light symptoms | 1449 |
| 5 | 2021-05-17 | 15-24yr | unvaccinated | hospitalisation | 1348 |
| 6 | 2021-05-17 | 15-24yr | unvaccinated | light symptoms | 989 |
| 7 | 2021-05-17 | 15-24yr | vaccinated | hospitalisation | 68 |
| 8 | 2021-05-17 | 15-24yr | vaccinated | light symptoms | 1466 |
| 9 | 2021-05-17 | 25-49yr | unvaccinated | hospitalisation | 1382 |
| 10 | 2021-05-17 | 25-49yr | unvaccinated | light symptoms | 1098 |

Showing 1 to 10 of 264 entries    Previous 1 2 3 4 5 … 27 Next

Download .csv

CTMA
Centre de Technologies Moléculaires Appliquées

UCLouvain

### Exercise n°3

Multiparametric simulator | 1) Datasets | 2) Simulations | 3) Visualization datasets | 4) Enrichment | 5) Visualization enrichment

#### Graph display of parameters

**Select the variable to be displayed in different colors (eg. variant)**

disease_outcome

**Select the variable to be displayed in different panels (eg. age_group)**

vaccination

**Do you want to filter with a third variable?**

- Yes
- No

Display graphs

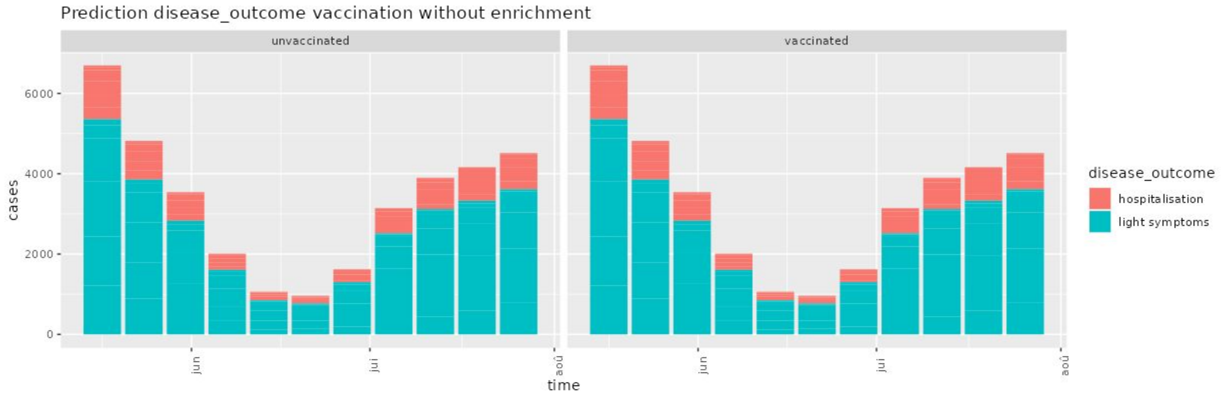


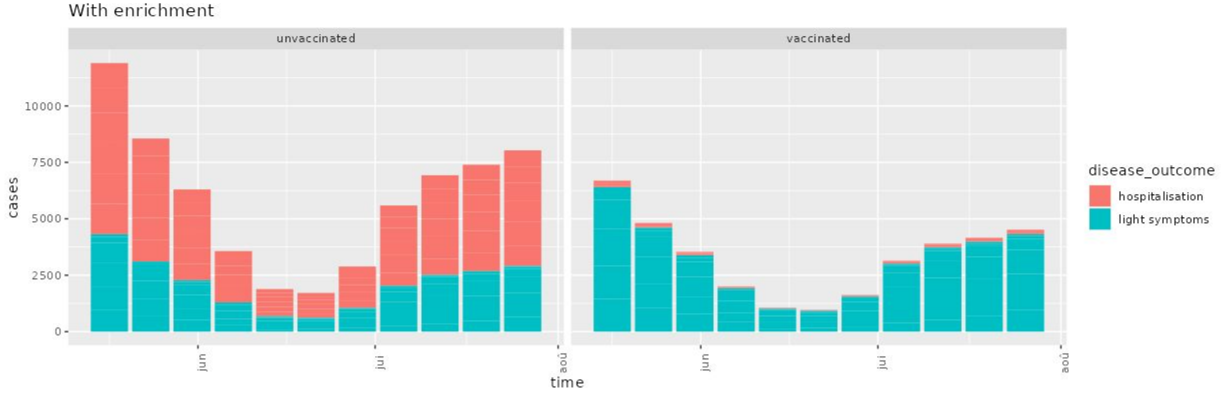
